# May mineral composition trigger or limit the protein content in soybean (*Glycine max* (L.) Merrill) seeds? Insights from a survey on 95 varieties cultivated in Brazil

**DOI:** 10.1101/2022.09.29.510200

**Authors:** Gabriel Sgarbiero Montanha, Lucas Coan Perez, Julia Rossatto Brandão, Rachel Ferraz de Camargo, Tiago Rodrigues Tavares, Eduardo de Almeida, Hudson Wallace Pereira de Carvalho

## Abstract

**Background and Aims:** Soybean (*Glycine max* (L.) Merrill) stands out as the major source of protein and oil for human and animal nutrition. Nevertheless, the increase in soybean yield has been accompanied by a reduction in its protein content in the last few decades. Since this might be influenced by the elemental composition of the seeds, we herein aimed at determining the profile of mineral nutrients and protein of 95 soybean varieties broadly cultivated in Brazil, the world’s biggest soybean producer and exporter, to identify possible nutritional triggers for the protein content.

**Methods:** Energy dispersive fluorescence spectroscopy (EDXRF) was employed to determine the concentration of macro, *i*.*e*., (K), phosphorus (P), sulphur (S), calcium (Ca), and micronutrients, *i*.*e*., iron (Fe), manganese (Mn), zinc (Zn), and copper (Cu). The protein content was evaluated in soybean seeds by the Dumas method. The correlational and clustering assessment between nutrients and protein were determined through both univariate and multivariate non-parametric tests.

**Results:** Both protein and nutrient concentrations are not homogeneous across soybean seed varieties, and a clear positive association between protein and sulphur (S), zinc (Zn), and manganese (Mn) concentrations were observed.

**Conclusion:** The recorded results suggest that sulphur (S), zinc (Zn), and manganese (Mn) are the limiting nutrients for higher protein content in soybean seeds.

## 1.0 INTRODUCTION

As the world’s population is expected to reach almost 10 billion people within the next few decades, there is a massive escalation in the demand for resources, especially nutritious food. At this juncture, the nourishment offer shall be increased by 70 % until 2050 to supply the foreseen population growth. Yet, it cannot be dissociated with environmental conservation (Food and Agriculture Organization of the United Nations 2009).

In this scenario, soybean (*Glycine max* (L.) Merril) stands out as a key crop, once its grain composition, which stores *ca*. 30-42 %wt. (weight basis) of protein without requiring nitrogen fertilisation (Dashiell 2005; Zong et al. 2017), one of the most expensive and least efficient agricultural inputs (Alves et al. 2003), as well as 18-22 %wt. of oil (Dashiell 2005; Zong et al. 2017)

According to data from the Food and Agriculture Organisation of the United Nations, the 2020 worldwide soybean production was *ca*. 13-fold larger than in 1961, yet within a *ca*. 5-fold larger area, which means that the production has expanded at a broader rate than the harvest areas (*ca*. 5 and 3 % year^-1^, respectively) in the last half-century (Food and Agriculture Organization of the United Nations 2021). It shows that soybean breeding programs combined with other technological advances, such as fertiliser broadcasting and mechanization, have been efficient in meeting the demand for this crop in the last decades. This skyrocketing yield is a major milestone for fulfilling the rising demand for protein and oil employed in human and animal nutrition. Additionally, in countries such as Brazil, 10 v. % of diesel is already composed of renewable biodiesel, this figure should be increased to 15 v. % in 2023 (Barros 2022).

Despite presenting a low concentration of sulphur-amino acids, *i*.*e*., methionine and cysteine, soybean is often reported as a high-quality vegetable protein source due to the presence of the nine essential amino acids, among them high contents of arginine, leucine, and lysine (*ca*. mass fraction of 9 %) (Young and Pellett 1994). Moreover, soybean seeds can also act as a source of mineral nutrients for human feeding, such as phosphorus (P), potassium (K), sulphur (S), calcium (Ca), manganese (Mn), iron (Fe), zinc (Zn), and copper (Cu).

In Brazil, soybean (*Glycine max* (L.) Merril) is currently the most cultivated crop, representing one of the main incomes in the country’s economic basket. In the 2021/2022 season, soybean global production reached the mark of 372.6 million Mg. From that, Brazil operates as the biggest player, producing 124.05 million Mg within 40.95 million hectares and average productivity of 3.029 tons/hectare. Brazil is also the world’s biggest soybean exporter. In 2021, the 74.1 million Mg of the commodity’s trading was worth US$ 26.1 billion, thus being the primary source of foreign currency in the Brazilian gross domestic product (GDP). Furthermore, exports of secondary products, such as soybean oil and meal amounted to 16.7 million and 1 million Mg, respectively, summarising US$6.6 billion (CONAB 2022).

In this context, although several strategies have been successfully explored for increasing the agricultural productivity of soybean crops, a major fraction did not focus on improving the quality of soybean seeds, such as the content or quality of soybean proteins, leading to a significant reduction on the protein content as a function of the higher outputs per unit of area (de Borja Reis et al. 2020; Umburanas et al. 2022). Overcoming this outline relies on understanding the physiological mechanisms that promote or hinder protein synthesis in soybeans seeds.

Therefore, the present study aimed at assessing the profile of mineral nutrients and protein of 95 soybean varieties broadly cultivated in Brazil in order to identify possible nutritional triggers for the protein content.

## 2.0 MATERIALS AND METHODS

### 2.1 Soybean varieties selection and sample preparation

Ninety-five (n=95) soybean varieties widely cultivated in the Brazilian southeast and midwest regions, main grain-producing areas in the country, were considered for this study. They were obtained from farmers, seed companies, and research institutions from Minas Gerais, São Paulo, and Goiás States, as detailed in Fig. 1 and the supplementary Table S1.

**Figure 1.**
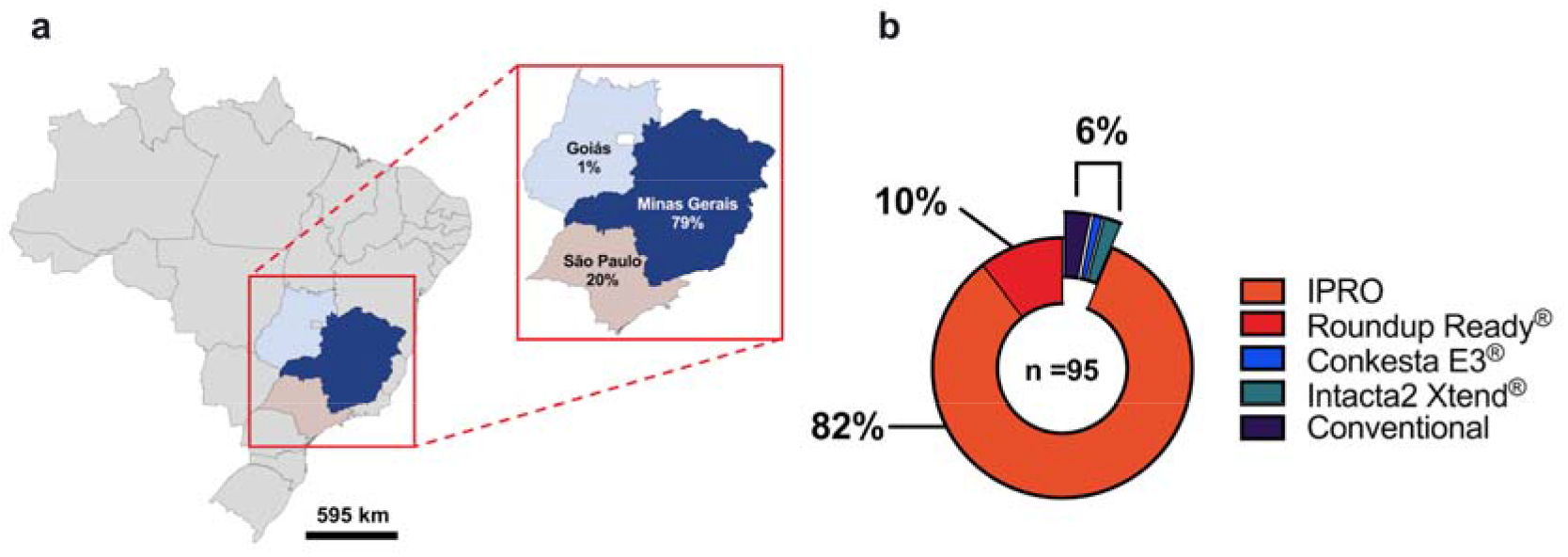
Schematic representation of the Brazilian States in which the 95 soybean varieties were acquired, whereas 95 and 25 % of the samples were collected from Minas Gerais and São Paulo, in southwest Brazil, and 1 % from Goiás, in the midwest (a). Profile of the varieties selected, revealing that the major fraction (96 %) consisted of glyphosate and caterpillars-resistant transgenic lines. Please check the supplementary Table S1 for a detailed description of all varieties herein employed (b).

The number of varieties selected, *i.e*., n = 95, was determined according to Equation 1 to reach the minimum sample size representing the 4567 soybean varieties (within a 95 % confidence level), which are listed at the National Registry of Cultivars from the Ministry of Agriculture, Livestock, and Supply (RNC-MAPA) on September 1, 2022 (Brazilian Ministry of Agriculture, 2022)

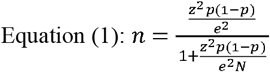

Where: N is the population size, *i.e*., 4567 soybean varieties listed at the National Registry of Cultivars from the Ministry of Agriculture, Livestock and Supply (RNC-MAPA) on September 1, 2022; e is a 10 % error margin; z is the score for a 95 % confidence interval, *i.e*., 1.96.

The soybean grains were ground using an electric blade grinder (Cadence, model MDR302, Brazil) for two cycles of 30 s each to obtain a homogeneous sample thereupon employed in all the subsequent analyses.

### 2.2 Elemental quantification

The elemental composition of the sampled soybeans was determined by Energy Dispersive X-ray Fluorescence Spectrometry (EDXRF, EDX-720, Shimadzu, Japan). 100 mg of the ground soybean were added into and X-ray sample cups (Spex SamplePrep no. 3577, USA) sealed with a 6-µm thick polypropylene film (VHG, FPPP25-R3 USA) at the bottom edge. After, the sample was gently pressed with a glass stick to prevent voids spaces. Subsequently, *ca*. 20 mg of the analytical grade boric acid was added over the sample and pressed with the glass stick again. Each sample was irradiated for 200 s by a 3-mm X-ray beam incoming from a Rh X-ray tube operating at 50 kV and with an auto-tunable current, so the detector dead-time does not exceed 30 % of the total. The X-ray spectra were collected by a Si(Li) detector. The analyses were carried out under vacuum using at least three independent replicates.

The concentration of P, S, K, Ca, Fe, and Mn was determined through the EDXRF fundamental parameter method, whereas Cu and Zn were determined using cellulose-based external calibration curves (Fig. S1), according to the procedure previously described ((Montanha et al., 2020)). In this latter case, both standards and samples analytes intensities were normalized by their respective background counting rates. The trueness of these methods was evaluated by using standard reference materials (NIST 1515, Apple Leaves) and soybean samples spiked with Zn standard solution at 1,000 mg L^-1^ (SpecSol® AAZN100V and AACU1000V, HNO_3_ 0.1-10 % v/v matrix) at 25 and 50 mg kg^-1^ of Cu and Zn, respectively. The recoveries are presented in Table S2.

### 2.3 Protein quantification

The quantification of protein in soybean seeds was carried out according to the Dumas official methodology from the Association of Official Analytical Chemists (AOAC, 1997). In this regard, the homogenized ground seed samples were analyzed using a Nitrogen/Protein analyzer (FP-528, Leco, USA). A 6.25 nitrogen-protein conversion factor was herein employed. The trueness of the method was determined using a soybean flour standard reference material (NIST 3234, USA), and the recovery was 101.4 %. All measurements were carried out using at least two independent replicates.

### 2.4 Statistical analysis

The normality of the data was assessed through the Shapiro-Wilk test at 0.05 level, as detailed in Table S3, and then classified into three groups as a function of the protein concentration range, *i.e*., ‘low protein’ (up to the 1^st^ quartile = 0-25 % of the data), ‘medium protein’ (between 1^st^ and 3^rd^ quartile = 26-75 % of the data), and ‘high protein’ (from the 3rd quartile upwards = 76-100 % of the data). The differences between each group were evaluated through Kruskal–Wallis one-way analysis of variance followed by Dunn’s multiple comparisons test at a 95 % confidence level (p < 0.05). Two-variable correlations were determined through Spearman’s rank correlation coefficient test at a 95 % confidence interval, followed by a K-means hierarchical clustering. Finally, Principal Component Analysis (PCA) was performed selecting the principal components with the largest eigenvalues able to explain at least 75 % of the total variance. All analyses were carried out using Prism (version 9.4.0, USA) and the packages ‘*stats*’ and ‘*ComplexHeatmap*’ of RStudio software (version 1.4.1106 “Tiger Daylily”, USA).

## 3.0 RESULTS

Figure 1a details the profile of 95 soybean varieties sampled in the States of Minas Gerais, São Paulo, and Goiás. It reveals that the major fraction of the samples proceeds from Minas Gerais (79 %) and São Paulo States (20 %), both in southwest Brazil, whereas only 1 % comes from the Goiás state, in the midwest region. These States were chosen because of an established professional network in these regions. The specific origin of each variety is detailed in Table S1.

Besides, one should notice that only 3.2 % (n=3) of the varieties were non-transgenic, *i.e*., whereas those containing transgenic lines, such as the Roundup Ready®, *i.e*., glyphosate-tolerant (Padgette et al., 1995) and the IPRO, *i.e*., resistant to both glyphosate and Lepidoptera caterpillars (Silva et al., 2019), encompass 92 % of the total.

Figure 2a presents the protein concentration found in soybean seeds from the 95 soybean varieties. It reveals that protein level ranges from 31.2 up to 39.8 % (percentage by mass basis), with average and median values of 35.3 and 35.5 %, respectively. It was thereby possible to classify them into ‘low’, ‘medium’, and ‘high’ protein groups as a function of the quartiles that encompass 0-25, 26-75 and 76-100 % of the data, respectively. The protein concentration obtained in each variety within these three groups are displayed in Fig. 2b, and their average values, *i.e*., 33.2, 35.6, and 37.7 % are shown in the boxplots shown in Fig. 2c.

**Figure 2.**
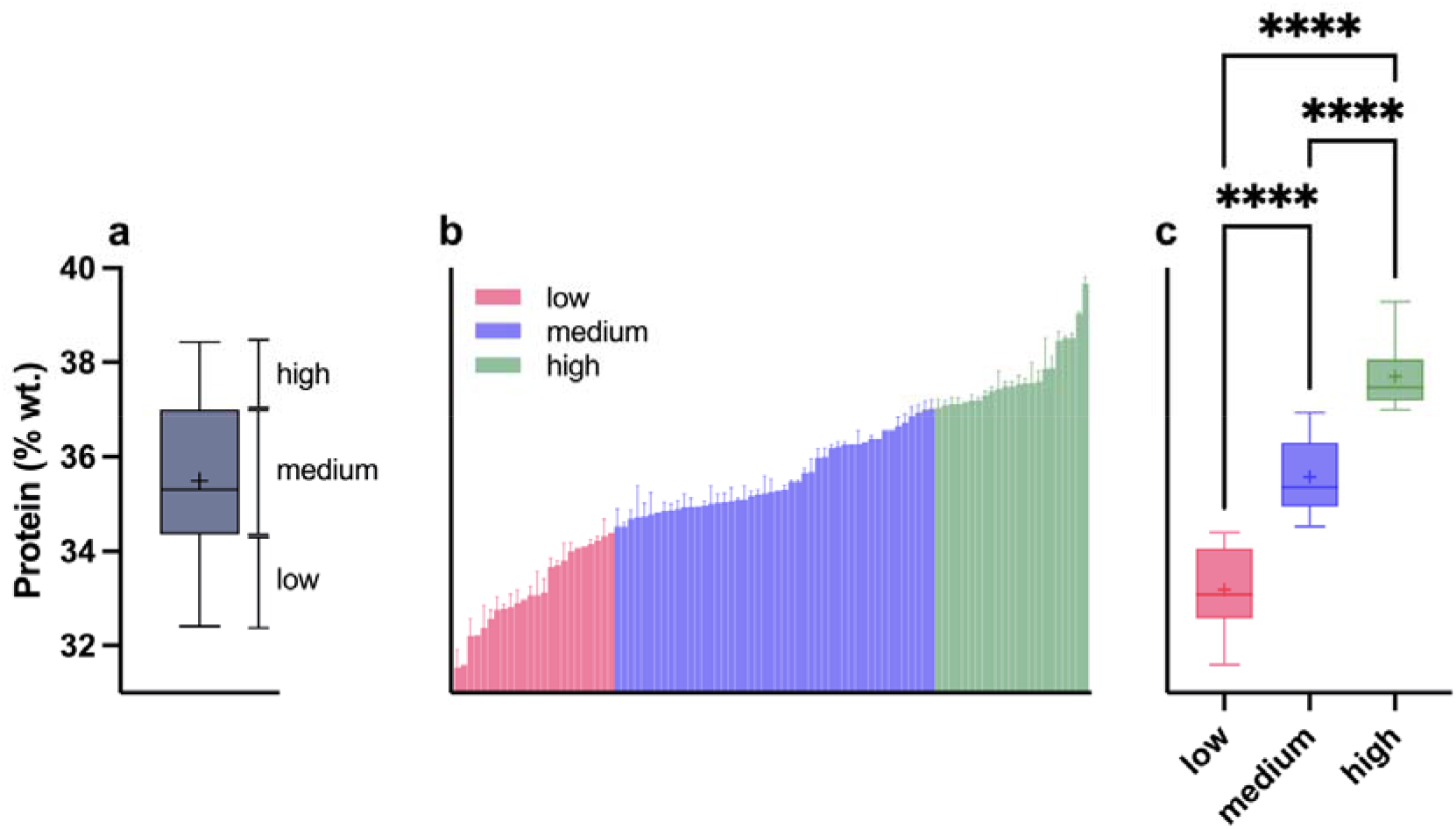
Boxplot of the protein concentration distribution found on the 95 soybean seed varieties, as well as their classification into low, medium, and high protein content groups, based on the quartile values (a). Ascending average protein concentration within the low, medium, and high protein content groups (b). Boxplot of the protein concentration distribution across low, medium, and high protein content groups (c). Data subjected to Kruskal–Wallis one-way analysis of variance followed by Dunn’s multiple comparisons test at 0.05 level. Data derived from at least two independent measurements. (+) mean values; (****) p <0.0001.

Figure 3 shows macro (*i.e*., K, P, S. and Ca; Fig. 3a) and micronutrients (*i.e*., Fe, Mn, Cu, and Zn; Fig. 3b) concentrations determined in the 95 soybean seed varieties via energy dispersive X-ray fluorescence spectroscopy (EDXRF). The concentration of each nutrient is significant different from each other, according to the Kruskal–Wallis one-way analysis of variance followed by Dunn’s multiple comparisons. Among the macronutrients, K presented the highest concentration values (ranging from *ca*. 16,000 up to *ca*. 20,000 mg kg^-1)^, followed by the content of P, S, and Ca, which presented average values of ca. 6,300, 3,300, and 2,200 mg kg^-1^, respectively. On the other hand, the micronutrient values decreased in the order Fe, Zn, Mn, and Cu, which showed average concentration values of 83, 35, 25, and 7 mg kg^-1^, respectively. The histograms shown in Fig. S2 detail the distribution of both protein and nutrient concentrations.

**Figure 3.**
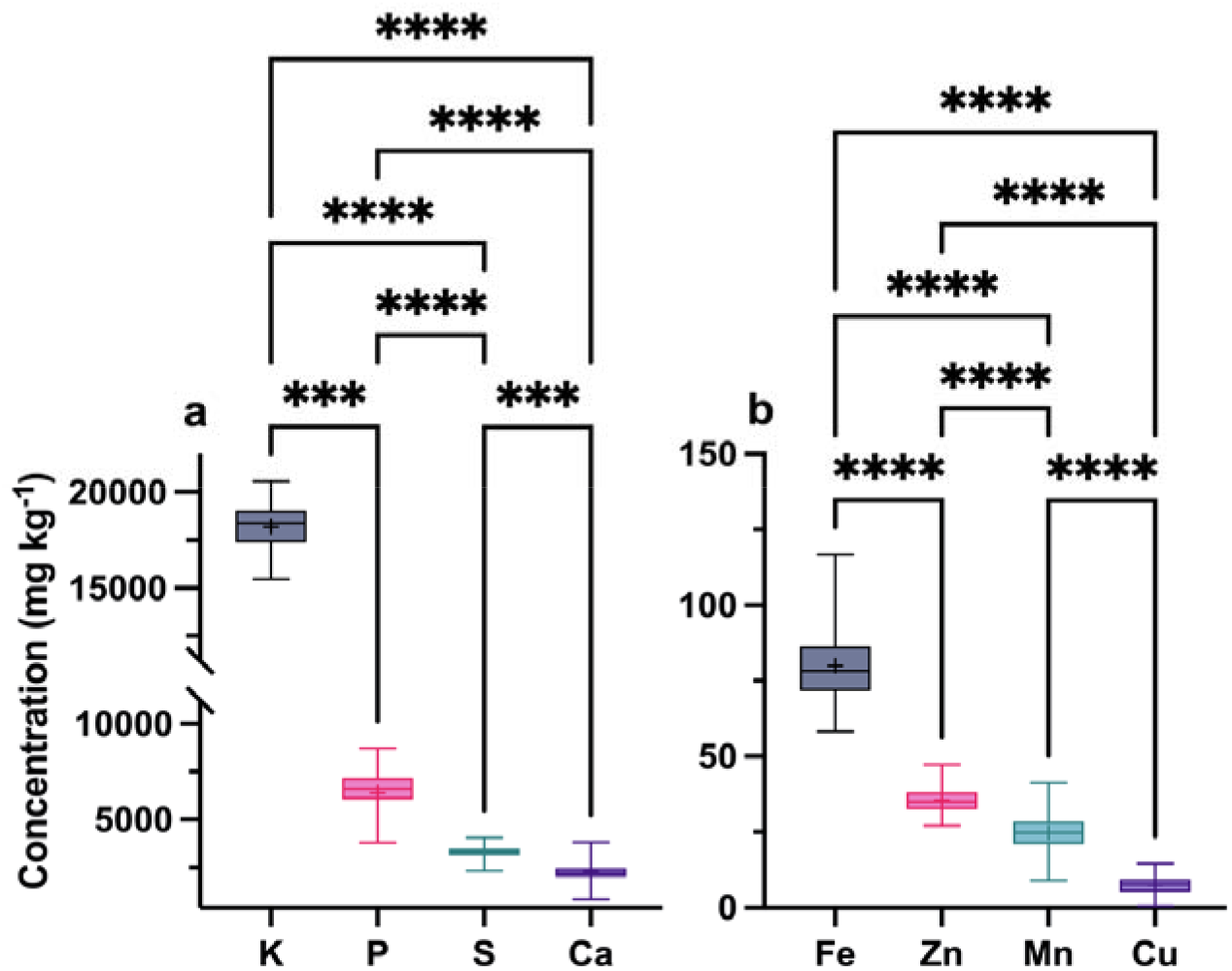
Boxplots of the concentration of macronutrients, *i.e*., K, P, S, and Ca (a), and micronutrients, *i.e*., Fe, Mn, Cu, and Zn (b) found on the 95 soybean seed varieties sampled at Minas Gerais, São Paulo, and Goiás States, Brazil. Data subjected to Kruskal–Wallis one-way analysis of variance followed by Dunn’s multiple comparisons test at 0.05 level. Measurements of elemental content were obtained from three independent replicates. Mean values were indicated with a “+”; Differences with p-value lower than 0.001 and 0.0001 were indicated with three and four asterisks.

The changes in the distribution of both macro and micronutrients concentrations as a function of the different protein content groups established through the quartile-based clustering is shown in Fig. 2a. Although the boxplots demonstrate a high dispersion of the data, it reveals a consistent drop in K and P mean concentrations as protein content increases, whereas an opposite trend is observed for S, Zn, Mn, and Cu concentrations, i.e., elemental contents increase together with protein contents. This trend is less pronounced for Zn, Mn and Cu than for S; however, one can note the first three elements showed concentrations statistically higher in samples with high protein contents than in samples with low and medium contents (Fig. 4). On the other hand, no clear changes were observed for Ca and Fe. The scatter plots shown in Fig. S3 itemizes these trends. Considering the outputs from Dunn’s *posthoc* test, it is thereby possible to state that both K, S, and Mn concentrations are more significantly affected by changes in the protein levels in the soybean seeds cultivated in the midwest and southeast Brazil (p <0.001).

**Figure 4.**
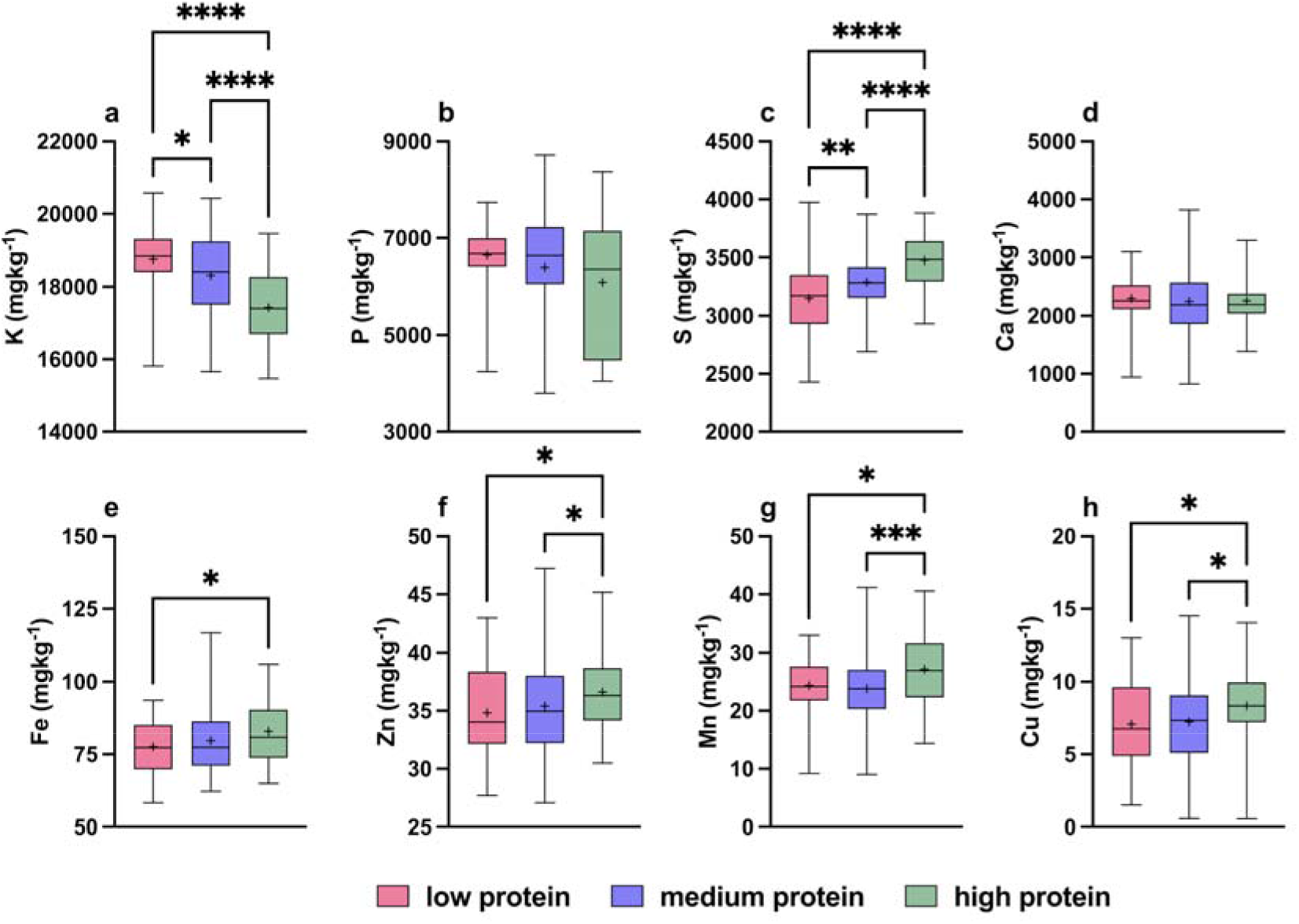
Boxplots of macronutrients, *i.e*., K, P, S, and Ca (a-d), and micronutrients, *i.e*., Fe, Zn, Mn, and Cu (e-h) as a function of the protein concentrations groups determined by the quartile-based clustering, *i.e*., ‘low’ (up to the first 25 % of the protein data range), ‘medium’ (protein range within 26-75 % of data), and ‘high’ (data upwards of 75 %) in the 95 soybean seed varieties sampled at Minas Gerais, São Paulo, and Goiás States, Brazil. Data subjected to Kruskal–Wallis one-way analysis of variance followed by Dunn’s multiple comparisons test at 0.05 level. Measurements of elemental content were obtained from three independent replicates. Mean values were indicated with a “+”; Differences with p-value lower than 0.1, 0.01, 0.001, and 0.0001 were indicated with one, two, three, and four asterisks.

In addition, the effects of sample location on both protein and nutrient concentration in the soybean surveyed were also assessed, as demonstrated in Fig. S4. It showed that the varieties sampled at Tupã, Iracemápolis, and Piracicaba (all cities from the São Paulo state) presented higher protein content, as well as higher Cu content compared to those from Minas Gerais state, *i.e*., the municipalities of Sacramento, Uberaba e Perdizes.

Finally, Fig. 5a displays the results of Spearman’s correlational analysis. The outputs unveil a stronger negative correlation (−0.5) between protein and K concentration, whereas milder ones were observed for P (−0.2) and Ca (−0.1). On the other hand, stronger positive correlation values were recorded for S (0.5) in relation to the weaker positive correlations observed for Zn, Cu, Fe (0.2), and Mn (0.1). Interestingly, the hierarchical k-means clustering presented the elements with negative correlation (S, K, and Ca) in the same group, while elements positively correlated with protein were allocated to the second one. This is corroborated by the principal component analysis presented in Fig. 5b, which clearly shows the loadings values of protein altogether with the ones of S and Mn, whereas P and K are displayed in the opposite direction.

**Figure 5.**
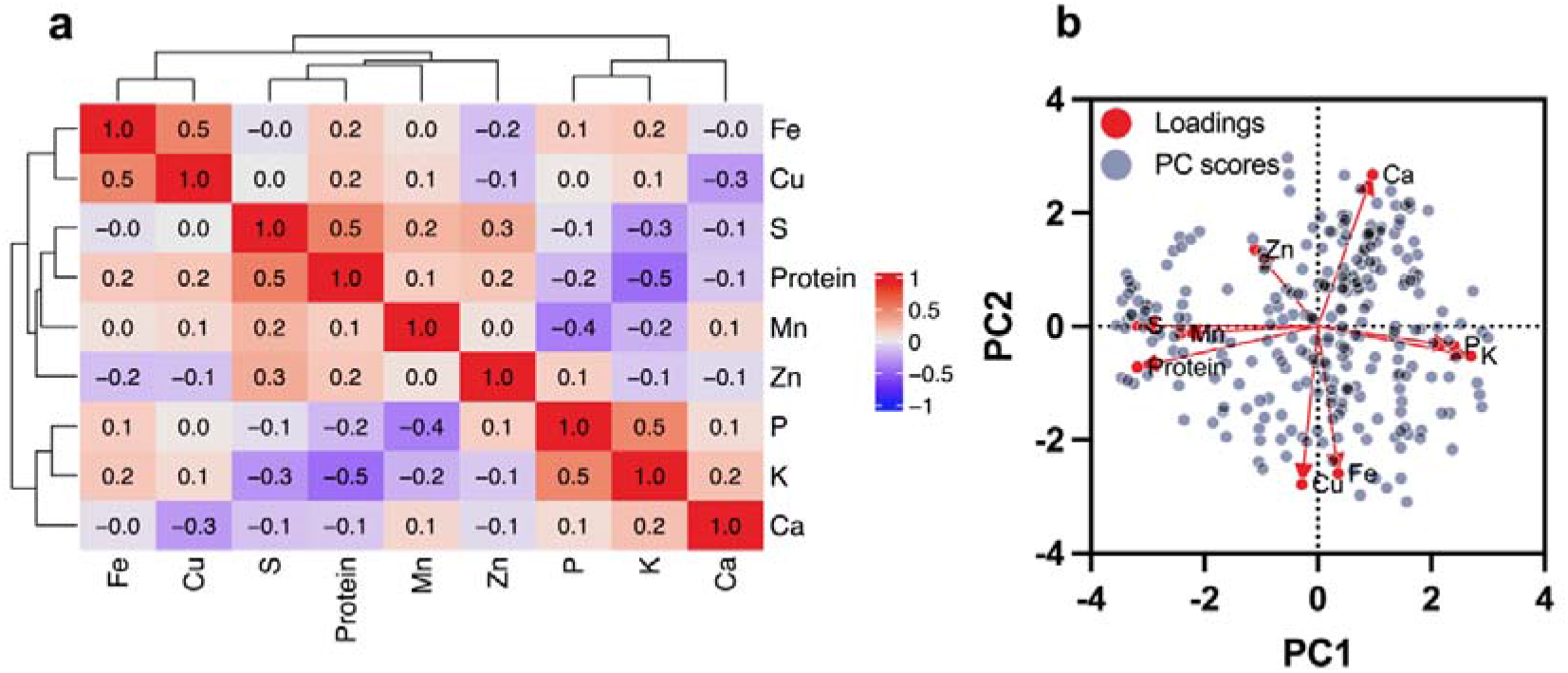
Heatmap of the Spearman’s rank correlation coefficient test at 95% confidence interval followed by K-means hierarchical clustering (a) and vector chart of the principal component analysis encompassing largest eigenvalues able to explain at least 75 % of the total variance of the protein, macronutrients, *i.e*., K, P, S, and Ca, and micronutrients, *i.e*., Fe, Zn, Mn, and Cu, concentration recorded in the 95 soybean seed varieties sampled at Minas Gerais, São Paulo, and the Goiás States, Brazil (b).

## 4.0 DISCUSSION

According to the Brazilian National Supply Company, Minas Gerais, São Paulo, and the Goiás States comprised *ca*. 19 % of the area used for soybean production in Brazil in the 2020/2021 harvest, (7,360,300 ha, an area larger than current Ireland’s territory), encompassing *ca*. 1/5 of Brazil’s total soybean output, *i.e*., 14,993,100 Mg, with higher productivity (3.7 Mg ha^-1^) compared to the national average (3.5 Mg ha^-1^) (CONAB, 2022).

One should notice that although productivity depends on several factors, the introduction of genetically modified (GM) strains, such as the glyphosate and caterpillars-tolerant technologies, *i.e*., Roundup-Ready® and IPRO (Padgette et al., 1995; Silva et al., 2019), features an enormous advantage for reaching higher outcomes. In this scenario, it has been estimated that GM varieties encompassed ca. 85 % of the total soybean cultivation area in Brazil during the 2011/2012 season (Homrich et al., 2012). This is highly consistent with the soybean seeds profile explored in this study, in which only 3 out of 95 varieties were conventional ones.

Nevertheless, several pieces of evidence suggest that higher productivity neither boosted the content nor quality of protein of soybean seeds, since a negative correlation between yield and the protein content has been often reported (Assefa et al., 2018; de Borja Reis et al., 2020; Mello Filho et al., 2004). A recent study conducted with 26 soybean cultivars released over the last 50 years in southern Brazil revealed that the yield has increased at a rate of 45.9 kg ha−1 year^-1^ whereas the protein content has decreased by 0.84 g kg−1 year^-1^, respectively (Umburanas et al., 2022). It means that the total grain weight yield is increasing at a rate of 21 % per decade, whereas the protein concentration is decreasing at a rate of ca. 1.1 % per decade. In 1960 the protein yield was *ca*. 0.72 Mg ha^-1^, considering the past protein concentration. Then, if the 1960 varieties were as productive as those of 2018, we should be harvesting 1.68 Mg of protein ha^-1^ instead of 1.59 Mg of protein ha^-1^.

Besides this downward trend in protein content, no significant changes in the amino acid composition were observed in a field trial exploring the soybean genotypes released between 1980 and 2014 in the United States (de Borja Reis et al., 2020). Thus, a change in amino acid profile would indicate an imbalance in the protein fractions, *i.e*., those from the albumin, globulin, glutelin, and prolamin families (Makeri et al., 2017), which has not been observed. It is important to highlight that most of the proteins in the soybean seeds are storage proteins ones (Derbyshire et al., 1976), *i.e*., a source of nitrogen to be used during seed germination and early-seedling development (Wakasa and Takaiwa, 2013). The glycinin and β-conglycinin, both salt-soluble proteins from the globulin family, encompass a major fraction of the storage proteins in soybean seeds (Murphy, 2008).

The present study shows that protein content in soybean seeds of the currently cultivated varieties is not homogeneous, ranging from 31.2 % up to 39.8 %, with an average value of 35.3 % (Fig. 2). It also varied across the cities in which the samples were collected (Fig. S4a). Similar numbers were found in soybean seeds cultivated in the United States (Assefa et al., 2019). Likewise, the concentration of both macro and micronutrients present in the grain has significantly spanned across the varieties (Fig. 3, Fig. S2).

As these figures are affected by environmental conditions and crop management practices (Assefa et al., 2019; de Borja Reis et al., 2020), one should expect that the protein content in soybean seeds might surely be influenced by their elemental composition. In this scenario, direct relationships were observed between protein content and the sulphur concentration (p = 0.5), as well as zinc, copper (p > 0.1-0.2), and manganese concentration in the probed soybean seeds (Fig. 4-5). Although relation does not necessarily imply causality, all four elements play crucial roles throughout the protein biosynthesis metabolic routes in plants.

Despite playing important roles in a myriad of metabolic processes (Borpatragohain et al., 2019; Q. Li et al., 2020; Narayan et al., 2022), sulphur encompasses both cysteine and methionine sulphur-amino acids residues in the primary structure of the glycinin, one of the most abundant fractions of globulin-like storage proteins in soybean seeds (Derbyshire et al., 1976; C. Li et al., 2019). In this scenario, sulphur-based foliar fertilisation has been reported to change the profile of soybean seed storage proteins (Ibañez et al., 2021).

Similarly, zinc has strong binding sites on glycinin (Nosworthy and Caldwell, 1987), and is a cofactor for the activity of methionine synthase in plants (Eckermann et al., 2000). Furthermore, zinc is also crucial for the proper functioning of key compounds in plant’s transcription, translation, antioxidant and catabolic process, *e.g*., RNA and DNA polymerases, ribosome associate proteins, CuZn superoxide dismutase (SOD1) and alcohol dehydrogenase (ADH), in which Zn-cysteine bounds are found (Broadley et al., 2007), and might then reflect the direct relationships observed between zinc and sulphur concentrations.

On the other hand, manganese has been associated with the phytin-rich globoid crystal within the proteins bodies (Buttrose, 1978; Madsen and Brinch-Pedersen, 2020), a specialised group of protein-storage vesicles found in all plant tissues, including seeds (Monma et al., 1992). Moreover, lower protein content was found in soybean seeds grown under limiting Mn concentrations (Carvalho et al., 2014; Wilson et al., 1982).

Together with the abovementioned publication, the present study provides sounding evidence that sulphur, zinc, copper, and manganese might act as nutritional triggers of proteins synthesis in soybean seeds, acting either as co-factors throughout the metabolic routes or even as the building blocks of the amino acid residues backbone. Or at least, they seem to be limiting factors to protein synthesis. As protein biosynthesis does not follow a linear progression in the seeds (Kambhampati et al., 2021), one should expect a spike in demands throughout particular timeframes. Therefore, a proper understanding of the protein accumulation profile in the soybean seeds might foster new fertilisation strategies able to enhance both their content and nutritional quality of storage proteins.

## 5.0 ACKNOWLEDGMENTS

The researchers involved in this study were funded by Stoller (FEALQ #104346) ICL América do Sul (FEALQ#104460), National Commission of Nuclear Energy (grant # 01341.001296/2021-11), and the São Paulo Research Foundation (20/07721-9 20/16670-9). H.W.P. Carvalho is the recipient of a research productivity fellowship from the Brazilian National Council for Scientific and Technological Development (CNPq) (grant # 306185/2020-2).

## 7.0 AUTHOR CONTRIBUTIONS

Gabriel Sgarbiero Montanha, Lucas Coan Perez, and Hudson Wallace Pereira de Carvalho contributed to the study’s conception and design. Material preparation, data collection and analysis were performed by Gabriel Sgarbiero Montanha, Lucas Coan Perez, Júlia Rosatto Brandão, Rachel Ferraz de Camargo, and Eduardo de Almeida. Data analysis was performed by Gabriel Sgarbiero Montanha and Tiago Rodrigues Tavares. The first draft of the manuscript was written by Gabriel Sgarbiero Montanha and Lucas Coan Perez and all authors commented on previous versions of the manuscript. All authors read and approved the final manuscript.

