## Supplementary Information for "May mineral composition trigger or limit the protein content in soybean (*Glycine max* (L.) Merrill) seeds? Insights from a survey on 95 varieties cultivated in Brazil"

^2^ Biology and Biotechnology Department “Charles Darwin”, Sapienza University of Rome. Via dei Sardi, 70, 00185 Rome, Italy.

^†^ These authors contributed equally to this work.

* To whom correspondence should be addressed:

Associate Professor Hudson Wallace Pereira de Carvalho.

| **Variety** | **Technology** | **Origin** | **Maturation Group** | **Variety** | **Technology** | **Origin** | **Maturation Group** | **Variety** | **Technology** | **Origin** | **Maturation Group** |
| --- | --- | --- | --- | --- | --- | --- | --- | --- | --- | --- | --- |
| RK7518 | IPRO | A | 7.5 | RK6719 | IPRO | B | 6.8 | 58I60RSF | IPRO | E | 6.8 |
| 74I77RSF | IPRO | A | 7.2 | ni | IPRO | B |  | 6968RSF | IPRO | F | 6.7 |
| DS7417 | IPRO | A | 7.4 | M7739 | IPRO | B | 7.7 | ni | IPRO | G |  |
| M7198 | IPRO | A | 7.1 | TEC7548 | IPRO | B | 7.5 | NS7007 | IPRO | H | 7.1 |
| TMG7067 | IPRO | A | 7.2 | CD2728 | IPRO | B | 7.2 | M5917 | IPRO | H | 5.9 |
| NS6906 | IPRO | A | 7 | LG60174 | IPRO | B | 7.4 | M6410 | IPRO | H | 6.4 |
| NS6990 | IPRO | A | ni | NS7667 | IPRO | B | 6 | NS8595 | IPRO | ni | 8.5 |
| LG60174 | IPRO | A | 7.4 | NS7709 | IPRO | B | 7 | NS8397 | IPRO | ni | 8.2 |
| 97R50 | IPRO | A | 7.5 | 74I77RSF | IPRO | B | 7.2 | SYN1687 | IPRO | ni | 8.7 |
| 97R70 | IPRO | A | ni | AS3730 | IPRO | B | 7.3 | NS8383 | RR | ni | 7.1 |
| NK7201 | IPRO | A | 7.2 | N57505 | IPRO | B | ni | 58I60RSF | IPRO | ni | 6.8 |
| RK6719 | IPRO | A | 6.8 | 73170RSF | IPRO | B | 7.3 | TMG7061 | IPRO | ni | 6.1 |
| TMG7067 | IPRO | A | ni | NK7201 | IPRO | C | 7.2 | 8473RSF | RR | ni | 7.4 |
| LG60162 | IPRO | A | 6.5 | 74I77RSF | IPRO | C | 7.2 | ni | IPRO | ni |  |
| 37179RSF | IPRO | A | ni | NS 6906 | IPRO | C | 7 | AS3590 | IPRO | ni | 5.9 |
| CD2728 | IPRO | A | 7.2 | 37179RSF | IPRO | C | ni | C2570 | RR | ni | 5.7 |
| C2618 | IPRO | A | ni | RK7518 | IPRO | C | 7.5 | DM57152RSF | IPRO | ni | 5.7 |
| 67168RSF | IPRO | A | ni | LG60162 | IPRO | C | 6.5 | C2626 | IPRO | ni | 6.1 |
| C2531E | Conventional | A | 5.3 | DS7417 | IPRO | C | 7.4 | ICS5619 | RR | ni | 5.9 |
| 96R29 | IPRO | A | 6.2 | NS6990 | IPRO | C | ni | ST592 | IPRO | ni | 5.9 |
| 8473RSF | RR | A | 7.4 | 98Y21 | IPRO | C | ni | FTR2155 | RR | ni | 5.8 |
| RK6316 | IPRO | A | 6.3 | DM73I75 | IPRO | C | 7.3 | NS7399 | IPRO | ni | 7.3 |
| CZ26B77 | IPRO | A | 6.7 | M6210 | IPRO | C | 6.2 | NS6990 | IPRO | ni | ni |
| B5560CE | CONKESTA E3 | A | 5.6 | TMG1180 | RR | D | 8 | NS7901 | RR | ni | 7.9 |
| NS6010 | IPRO | A | 6 | SN75172 | IPRO | D | ni | NS7780 | IPRO | ni | 7.8 |
| M6620I2X | INTACTA2 XTEND | A | 6.6 | ST797 | IPRO | D | 7.9 | NS6906 | IPRO | ni | 7 |
| NK6356 | IPRO | A | 6.3 | FPS1859 | RR | D | 5.5 | 83HO113 TP | IPRO | ni | 8.3 |
| AS3707I2X | INTACTA2 XTEND | A | 7 | 8473RSF | RR | D | 7.4 | CZ 48B32 | IPRO | ni | 8.3 |
| 95R95 | IPRO | B | ni | M8372 | IPRO | D | 8.3 | CZ 58B28 | IPRO | ni | 8.2 |
| DS7417 | IPRO | B | 7.4 | BRS232 | Conventional | E | 6.9 | NS7700 | IPRO | ni | 7.7 |
| ST777 | IPRO | B | 7.7 | BRS284 | Conventional | E | 6.3 | NS7790 | IPRO | ni | 7.7 |
| RK7518 | IPRO | B | 7.5 | 55157 RSF | IPRO | E | 5.5 |  |  |  |  |
| A = Perdizes (-19.343, -47.296); B = Sacramento (-19.862, -47.451); C = Uberaba (-19.747, -47.938); D = Tupã (-21.934, -50.519); E = Iracemápolis (-22.583, -47.523); F = Jumirim (-23.088, -47.787); G = Rio Verde (-17.792, -50.919; H = Piracicaba (-22.734, -47.648); ni - not informed by the suppliers | | | | | | | | | | | |

**Table S1.** Description of technology, origin and maturation group from the the 95 soybean seed varieties sampled at Minas Gerais, São Paulo, and Goiás states, Brazil.

| **Analyte** | **Determined (mg kg^-1^)** | | | **Reference values (mg kg^-1^)** | | **Recovery (%)** | | | |
| --- | --- | --- | --- | --- | --- | --- | --- | --- | --- |
| P | 1773.96 | ± | 64.10 | | 1593.00^*^ | | 114.21 | ± | 4.0 |
| S | 1754.01 | ± | 38.86 | | 1800.00^*^ | | 87.31 | ± | 8.2 |
| K | 15061.41 | ± | 173.30 | | 16080.00^*^ | | 94.43 | ± | 1.1 |
| Ca | 13112.25 | ± | 115.44 | | 15250.00^*^ | | 86.52 | ± | 0.8 |
| Mn | 50.22 | ± | 3.22 | | 54.10^*^ | | 88.62 | ± | 6.0 |
| Fe | 78.07 | ± | 0.39 | | 82.70^*^ | | 94.74 | ± | 0.5 |
| Cu | 24.57 | | | 24.00^**^ | | 101.51 | | | |
| Zn | 55.34 | | | 50.25^**^ | | 101.89 | | | |
| ^*^ SRM^Ⓡ^ NIST1515 (Apple Leaves) | | | | | | | | | |
| ^**^ spikes from 1000 µg mL^-1^ standard solutions on soybean seed samples | | | | | | | | | |

**Table S2.** Determined and reference concentration and their respective recovery values found P, S, K, Ca, Mn, Fe, Cu, and Zn determined through energy-dispersive X-ray fluorescence spectroscopy. The reference values consisted of NIST1515 (Apple Leaves) standard reference material or soybean samples spiked with Cu and Zn standard solutions.

| **Analyte** | **Protein level group** | **N** | **W** | **p-value** | **Normality** |
| --- | --- | --- | --- | --- | --- |
| Protein | low | 72 | 0.9716 | 0.1026 | Accept |
|  | medium | 144 | 0.942 | <0.0001 | Reject |
|  | high | 69 | 0.8465 | <0.0001 | Reject |
| K | low | 72 | 0.9257 | 0.0004 | Reject |
|  | medium | 144 | 0.9718 | 0.0045 | Reject |
|  | high | 69 | 0.9673 | 0.0673 | Accept |
| P | low | 72 | 0.8245 | <0.0001 | Reject |
|  | medium | 144 | 0.8967 | <0.0001 | Reject |
|  | high | 69 | 0.9099 | 0.0001 | Reject |
| S | low | 72 | 0.971 | 0.0947 | Accept |
|  | medium | 136 | 0.9941 | 0.8541 | Accept |
|  | high | 69 | 0.9751 | 0.1831 | Accept |
| Ca | low | 72 | 0.9375 | 0.0014 | Reject |
|  | medium | 144 | 0.9692 | 0.0025 | Reject |
|  | high | 69 | 0.9367 | 0.0017 | Reject |
| Fe | low | 69 | 0.9694 | 0.0886 | Accept |
|  | medium | 139 | 0.9421 | <0.0001 | Reject |
|  | high | 66 | 0.9629 | 0.0456 | Reject |
| Zn | low | 72 | 0.9634 | 0.0343 | Reject |
|  | medium | 143 | 0.9674 | 0.0017 | Reject |
|  | high | 69 | 0.9865 | 0.6656 | Accept |
| Mn | low | 72 | 0.9688 | 0.0711 | Accept |
|  | medium | 144 | 0.9955 | 0.9411 | Accept |
|  | high | 69 | 0.9831 | 0.4778 | Accept |
| Cu | low | 72 | 0.9682 | 0.0651 | Accept |
|  | medium | 144 | 0.9923 | 0.6318 | Accept |
|  | high | 69 | 0.9603 | 0.0276 | Reject |

**Table S3**. Output from Shapiro-Wilk’s normality test at 95% interval of confidence for the protein, macronutrients, *i.e.*, K, P, S, and Ca, and micronutrients, *i.e.*, Fe, Zn, Mn, and Cu as a function of the protein concentrations groups determined by the quartile-based clustering, *i.e.*, 'low' (up to the first 25 % of the protein data range), 'medium' (protein range within 26-75 % of data), and 'high' (data upwards of 75 %) in the 95 soybean seed varieties sampled at Minas Gerais, São Paulo, and Goiás States, Brazil.

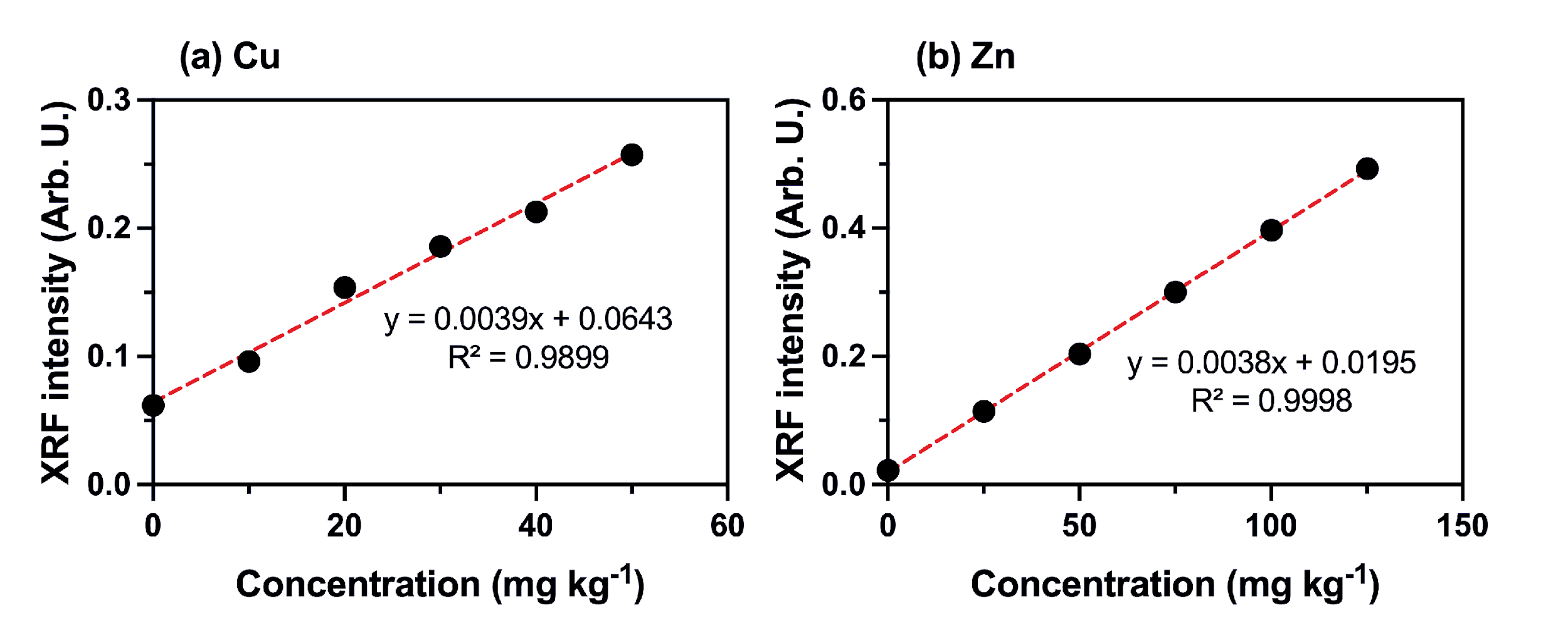

**Figure S1**. External calibration curve for X-ray fluorescence quantitative method, where the concentration of Cu and Zn in cellulose pellet standards (mg kg^-1^) is correlated with the respective normalized XRF intensities of standards.

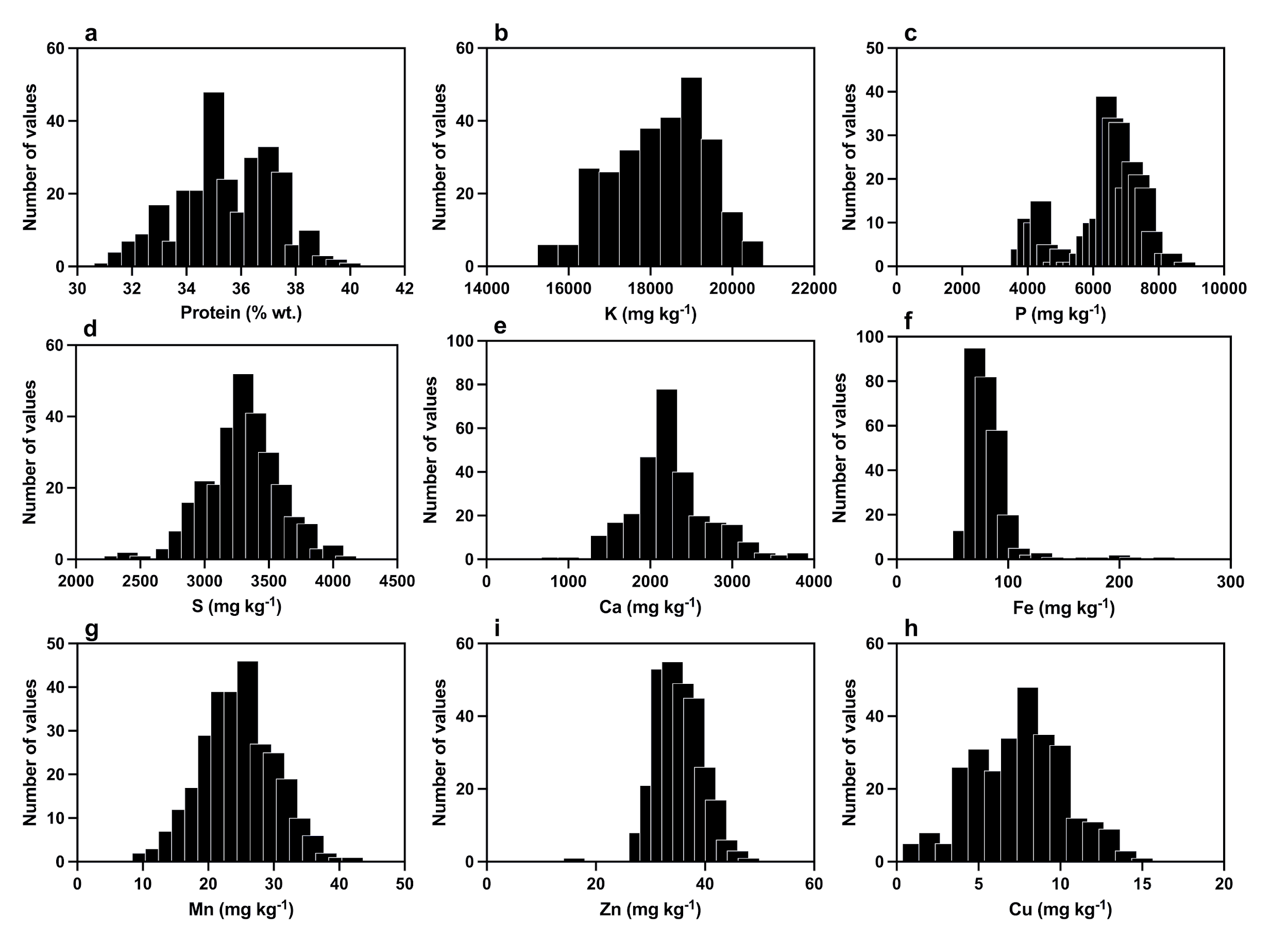

**Figure S2.** Histograms of the concentration of protein (a), macronutrients, *i.e.*, K, P, S, and Ca (b-e), and micronutrients, *i.e.*, Fe, Mn, Cu, and Zn (f-h) found on the 95 soybean seed varieties sampled at Minas Gerais, São Paulo, and Goiás States, Brazil.

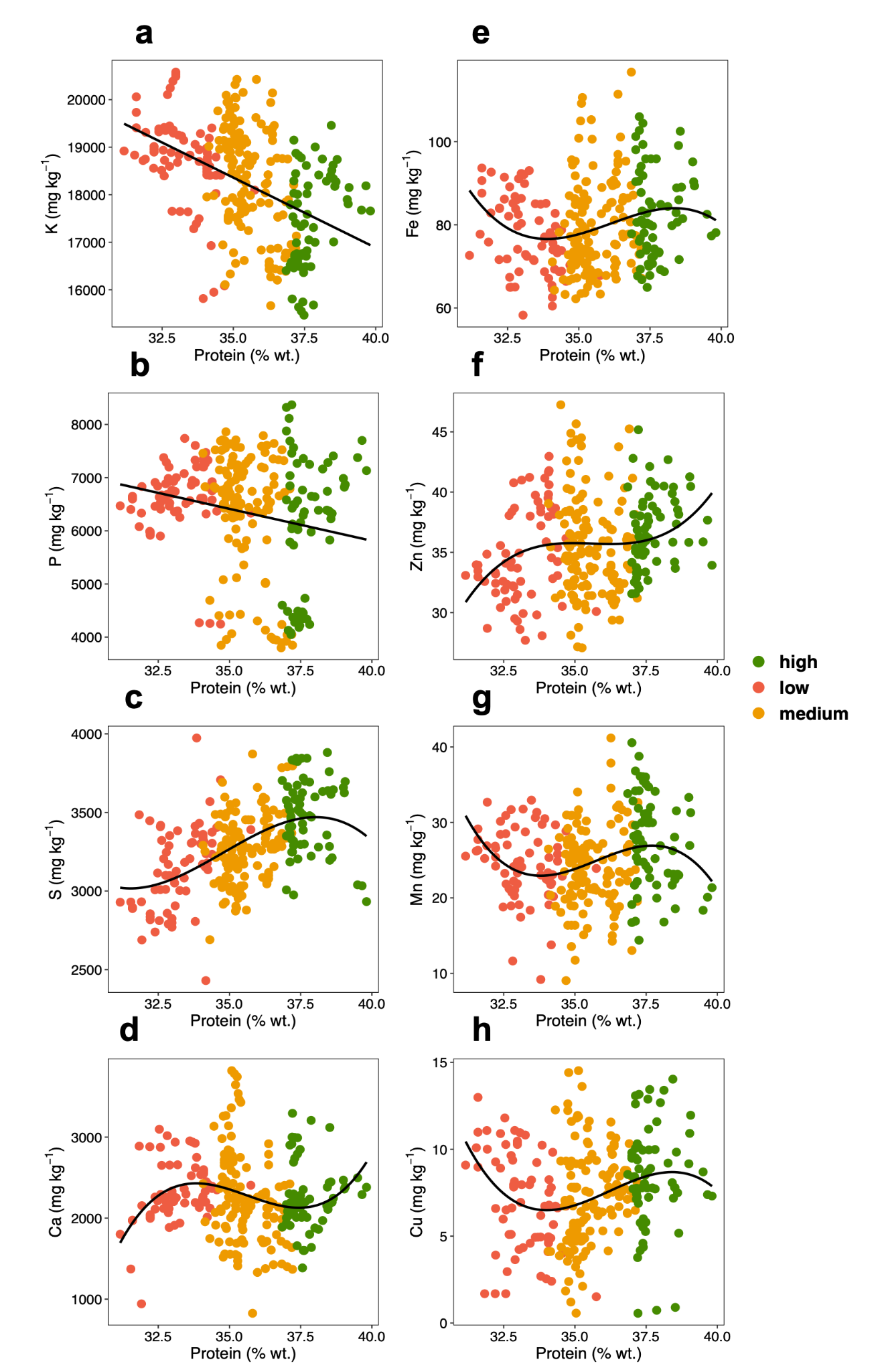

**Figure S3.** Scatter plots of macronutrients, *i.e.*, K, P, S, and Ca (a-d), and micronutrients, *i.e.*, Fe, Zn, Mn, and Cu (e-h) as a function of the protein content in the 95 soybean seed varieties sampled at Minas Gerais, São Paulo, and Goiás states, Brazil. Different colours indicate the protein concentrations groups determined by the quartile-based clustering, *i.e.*, 'low' (up to very first 25 % of the protein data range, in red), 'medium' (protein range within 26-75 % of data, in orange), and 'high' (upwards 75 % of the data, in green) in the 95 soybean seed varieties sampled at Minas Gerais, São Paulo, and Goiás states, Brazil. Data subjected to Kruskal–Wallis one-way analysis of variance. Data subjected to a 3^rd^ order polynomial regression.

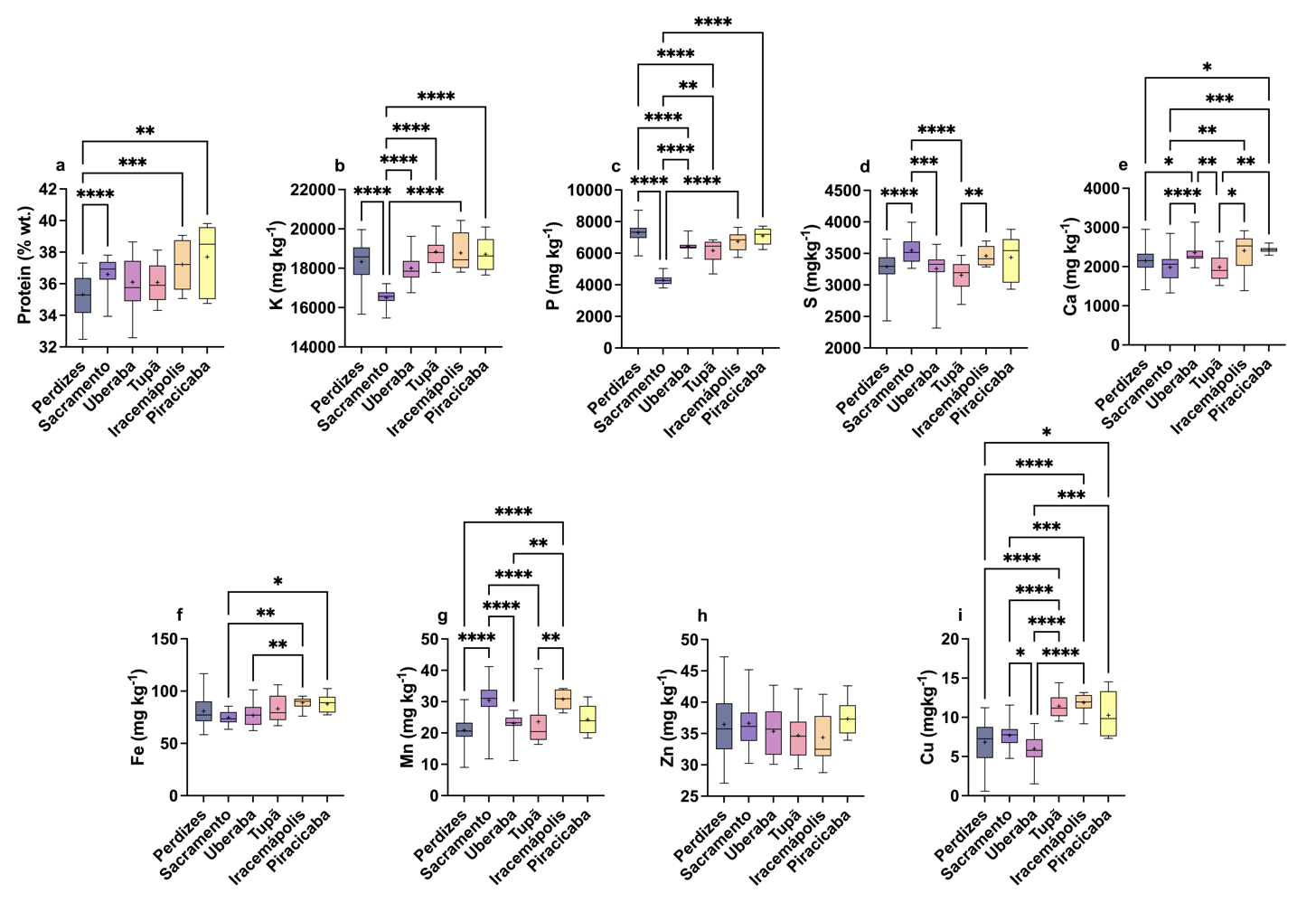

**Figure S4.** Boxplots of protein (a), macronutrients, *i.e.*, K, P, S, and Ca (b-e), and micronutrients, *i.e.*, Fe, Zn, Mn, and Cu (f-i) concentrations found in the cities from Minas Gerais and São Paulo, Brazil. Only the cities in which more than 3 varieties were sampled are herein shown. Data subjected to Kruskal–Wallis one-way analysis of variance followed by Dunn's multiple comparisons test at 0.05 level. Measurements of elemental content were obtained from three independent replicates. Mean values were indicated with a “+”; Differences with p-value lower than 0.1, 0.01, 0.001, and 0.0001 were indicated with one, two, three, and four asterisks.
